# MECOM permits pancreatic acinar cell dedifferentiation avoiding cell death under stress conditions

**DOI:** 10.1101/2020.08.21.260539

**Authors:** Elyne Backx, Elke Wauters, Jonathan Baldan, Mathias Van Bulck, Ellis Michiels, Yves Heremans, Diedert Luc De Paep, Mineo Kurokawa, Susumu Goyama, Luc Bouwens, Patrick Jacquemin, Isabelle Houbracken, Ilse Rooman

## Abstract

Maintenance of the pancreatic acinar cell phenotype suppresses tumor formation. Hence, repetitive acute or chronic pancreatitis, stress conditions in which the acinar cells dedifferentiate, predispose for cancer formation in the pancreas. Dedifferentiated acinar cells acquire a large panel of duct cell-specific markers. However, it remains unclear to what extent dedifferentiated acini differ from native duct cells and which genes are uniquely regulating acinar cell dedifferentiation. Moreover, most studies have been performed in mice since the availability of human cells is scarce.

Here, we applied a non-genetic lineage tracing method of human pancreatic exocrine acinar and duct cells that allowed cell-type-specific gene expression profiling by RNA sequencing. Subsequent to this discovery analysis, one transcription factor that was unique for dedifferentiated acinar cells was functionally characterized.

RNA sequencing analysis showed that human dedifferentiated acinar cells expressed genes in ‘Pathways of cancer’ with prominence of *MECOM (EVI-1), a* transcription factor that was not expressed by duct cells. During mouse embryonic development, pre-acinar cells also transiently expressed MECOM and in the adult mouse pancreas, MECOM was re-expressed when mice were subjected to acute and chronic pancreatitis, conditions in which acinar cells dedifferentiate. In human cells and in mice, MECOM expression correlated with and was directly regulated by SOX9. Mouse acinar cells that, by genetic manipulation, lose the ability to upregulate MECOM showed impaired cell adhesion, more prominent acinar cell death and suppressed acinar cell dedifferentiation by limited ERK signaling.

In conclusion, we transcriptionally profiled the two major human pancreatic exocrine cell types, acinar and duct cells, during experimental stress conditions. We provide insights that in dedifferentiated acinar cells, cancer pathways are upregulated in which MECOM is a critical regulator that suppresses acinar cell death by permitting cellular dedifferentiation.

## INTRODUCTION

Few cancers have such poor survival rates as pancreatic ductal adenocarcinoma (PDAC) (1). Understanding the molecular mechanisms that are at play in early-stage cancer formation is important to improve disease outcomes, but clinical samples of early disease in this case are rare. Hence, genetically modified mouse models are pivotal to provide us insight into the contribution of the different pancreatic cell types to tumor formation (2–4). Accumulating evidence points to acinar cells at origin of pancreatic tumor development (5–10). In order to become susceptible to tumorigenesis, acinar cells have to lose their differentiated phenotype to some extent. Consequently, tumorigenesis is restrained by mechanisms that control acinar cell differentiation (5–10). Loss of differentiation can result from repetitive acute or chronic pancreatitis whereby dedifferentiated acinar cells acquire a panel of markers that are typical for (embryonic) duct cells (11–13). Notably, duct cells can also give rise to tumors but likely under different conditions and through other mechanisms (14–16).

To date, knowledge is lacking on the mechanisms that govern the duct-like dedifferentiated acinar cell phenotype versus the ‘native’ duct cell phenotype, the latter being more resistant to stress- or inflammation-associated tumorigenesis. Previously, we established *in vitro* experimental models that recapitulate human (12) and rodent (13) acinar dedifferentiation. Specifically, upon suspension culture acinar cells lose their acinar-specific markers while gaining (embryonic) duct-like features, thereby mimicking chronic pancreatitis (12, 13). Here, we investigated the differential gene expression in adult human dedifferentiated acinar cells compared to native duct cells from the same cell preparations. The transcription factor *MECOM (EVI-1)* was found to be uniquely re-expressed-since embryonic development-in dedifferentiated acinar cells.

Although *MECOM* transcription factor is a known oncogene with a role in apoptosis (17–19) and pancreatic tumor formation (20, 21), its function in acinar cell (de)differentiation was unknown. We analyzed its role in the context of acinar cell dedifferentiation by caerulein-induced pancreatitis modeling in transgenic *Evi-1* knock-out mice. We provide mechanistic insights that mouse acinar cells are dependent on MECOM expression to prevent cell death, allowing them to acquire a dedifferentiated cell state.

## MATERIALS AND METHODS

### Human donor material and cell culture

Human healthy donor pancreatic exocrine cell preparations as residual material from a pancreatic islet donor program were processed by the Beta Cell Bank of the JDRF Center for Beta Cell Therapy in Diabetes (Brussels, Belgium), affiliated to the Eurotransplant Foundation (Leiden, The Netherlands). Consent for the use of residual donor material for research was obtained according to the legislation in the country of organ procurement. This project was approved by the Medical Ethical Committee of UZ Brussel - Vrije Universiteit Brussel (B.U.N. 143201732260). Human exocrine cells were cultured as previously described (11, 12).

### Mouse strains and experiments

ElaCreERT mice (22) and ElaCreERT;Sox9^f/f^ (23) originally from D. Stoffers (UPenn, USA) were kindly provided by P. Jacquemin (UCL, Belgium). ElaCreERT mice were crossed with EVI-1exon4 loxP mice (*Mecom*^f/f^) (24) (kindly provided by M. Kurokawa, UPenn), resulting in ElaCreERT;*Mecom^f^*^/f^ mice (Mecom KO). Six to eight-weeks-old female and male mice received tamoxifen and (Z)-4-Hydroxytamoxifen (all from Sigma-Aldrich, St-Louis, MO, USA) and were treated with 125µg/kg body weight caerulein (Invitrogen Waltham, Massachusetts, USA) according to (23). Mouse acinar cell isolation was performed according to a previously published protocol (13). Mice were not randomized but further processing and analyses of samples were performed blinded. Embryonic mouse tissue was isolated from ElaCreERT mice at E16.5 timepoint of embryonic development. All animal experiments were approved by the Ethical Committee for Animal Testing at the Vrije Universiteit Brussel (#17-637-1).

### Fluorescent activated cell sorting (FACS)

FITC-conjugated UEA-1 (*Ulex Europaeus* Agglutinin-1, Sigma-Aldrich) lectin labeling of acinar cells within the human exocrine cell fraction (isolated from healthy donor pancreata) was performed as in (12). At day 4 of suspension culture, cell clusters were dissociated following the protocol of (11). Analysis and cell sorting were performed on a BD FACSAria (BD Biosciences, Franklin Lakes, NJ, USA). Viable, single cells were gated based on forward and side scatter.

### Cell lines and culture

The mouse partially differentiated acinar cell line 266-6® CRL-2151 was purchased from American Type Culture Collection (ATCC, Manassas, Virginia, USA). Cells were maintained in DMEM + GlutaMax medium, supplemented with 10% heat-inactivated fetal bovine serum and 1% penicillin-streptomycin (all from Gibco) in a humidified atmosphere at 37°C and 5% CO_2_. All cell lines tested negative for mycoplasma contamination. Absolute cell number was determined by seeding 50 000 cells per well of a 6-well plate on day 0 and counted 96 h after seeding. CellTiter Glo assay (Promega, Madison, Wisconsin, USA) was carried out according to manufacturer’s instructions. In short, cells were seeded *in triplo* in opaque 96-well plates and cell viability was measured 96h later using the GloMax Discover Microplate reader (Promega). Luciferase reporter assay (Promega) was carried out according to the manufacturer’s instructions. Briefly, 1 000 cells were seeded *in triplo* in opaque 96-well plates and transfected 24 h later with a total of 200 ng DNA (pcDNA3.1-Sox9 only, pGL3-E1Bp-MECenh (a kind gift by R. Cullum, BC Cancer Research Centre, Vancouver, Canada) only or co-transfection with both). Luciferase activity was measured 48 h later using the GloMax ® Discover Microplate reader.

### CRISPR plasmids

Guide RNAs (gRNAs) for MECOM CRISPR knockdown were designed using the MIT CRISPR tool (https://zlab.bio/guide-design-resources). The top three gRNAs with the least predicted off-target activity were selected and ordered as oligos and cloned into the lentiCRISPRv2-PURO vector, a kind gift from B. Springer (Addgene plasmid #98290; https://www.addgene.org/98290/), following the manufacturer’s instructions.

### Lentiviral transduction and selection

266-6 cells were seeded at a density of 50 000 cells per well of a 6-well plate. 24 h later, cells were transduced with lentiviral supernatants supplemented with protamine sulphate (10 µg/mL, Sigma-Aldrich) for 48 h. After splitting, cells were selected for at least 1 week in 0,4 µg/mL puromycin (Invitrogen) before any further analysis was performed. Human exocrine cells were transduced with lentiviral shRNA-Evi1 vectors (MOI 10) on the day of isolation for 16 h at a density of 200 000 cells in Advanced RPMI supplemented with 5% heat-inactivated FBS, 1% GlutaMax and 1% Pen/Strep supplemented with protamine sulphate (10 µg/mL, Sigma-Aldrich). Cells were then washed with PBS and further cultured for 4 to 7 days in Advanced RPMI supplemented with 5% heat-inactivated FBS, 1% GlutaMax and 1% Pen/Strep.

### Immunostainings

Pancreas tissue and cell pellets were fixed over night at room temperature in 10% neutral-buffered formalin (KliniPath, Olen, Belgium). Next, samples were washed twice with PBS, dehydrated and embedded in paraffin. Sections of 4 µm thickness were cut. Automated stainings were performed on the Leica Bond RX™. Primary antibodies used are: anti-EVI-1 (1/1000, C50E12, Cell Signaling Technology, Danvers, MA, USA), anti-cleaved caspase 3 (1/200, 9661, Cell Signaling Technology), anti-CD3 (1/50, A0452, Agilent), anti-Krt19 (1/100, TromaIII, Developmental Studies Hybridoma Bank, Iowa City, IA, USA), anti-F4/80 (1/100, APC clone BM8.1, Sigma-Aldrich), anti-CD142 (1/100, AF2419, Bio-techne, Minneapolis, MN, USA), anti-Krt19 (1/20, M0888, Agilent, Santa-Clara, CA, USA), anti-CD19 (1/500, ab245235, Abcam), anti-collagen IV (1/500, 1340-01, Southern Biotechnology associates) Trichrome Masson’s staining was performed automated at the Pathology Department of UZ Brussel, Brussels, Belgium. DNA staining was performed with DAPI (Agilent) or Hoechst.

### RNA in situ hybridization

RNA in situ hybridization (RISH) was performed on formalin-fixed, paraffin-embedded tissue sections, according to the manufacturer’s protocols for manual RNAscope^®^ 2.5 HD Assay—RED (Bio-techne). Images were acquired using an EVOS FL1 auto digital microscope (Thermo Fisher Scientific). Immunostainings and average mRNA dots (RISH) were determined using HALO® image analysis platform (Indica Labs) or FIJI® imaging software.

### Quantitative Reverse Transcription Polymerase Chain Reaction (qRT-PCR)

Total RNA was isolated from cells using NucleoSpin RNA isolation kit (Macherey-Nagel, Düren, Germany) or GenElute Mammalian Total RNA Miniprep kit (Sigma-Aldrich). RNA concentration was measured by NanoDrop2000 (ThermoFisher). cDNA was prepared using the GoScript Reverse Transcription System (Invitrogen). qPCR was performed using FastSYBRGreen 5x MasterMix on a QuantStudio 6 (Invitrogen). Analysis was done using the ΔΔCt method using HPRT as housekeeping gene.

### Immunoblotting

Total protein lysates were extracted with RIPA buffer (150mM NaCl, 1.0% Triton X-100, 0.5% sodium deoxycholate, 0.1% SDS and 50mM Tris pH 8.0) supplemented with protease inhibitor cocktail and phosphatase inhibitor cocktail (all from Sigma-Aldrich). Protein concentration was measured using Bradford assay (Bio-Rad, Hercules, CA, USA). Protein extracts were mixed with 5x Laemmli buffer (4% SDS, 20% glycerol, 10% ß-mercaptoethanol, 0,004% bromophenol blue and 0,125M Tris-HCl pH 6.8) and reduced by boiling for 5 min. at 95°C. Samples were loaded on Bolt 4-12% gels (Invitrogen) and transferred onto PVDF membrane (Invitrogen). After 1 h blocking in 5% milk or 5% BSA in TBS-T, blots were incubated with 5mL Revert Total Protein Stain (Li-Cor Biosciences, Lincoln, NE, USA) for 5 min., followed by 3 washes with TBS-T and imaged on the LiCor Odyssey Fc Imaging System for normalization of protein expression. Then, primary antibodies were incubated at 4°C in 3%BSA in TBS-T.

Primary antibodies used were: anti-EVI1 (C50E12, Cell Signaling Technology, Danvers, MA, USA, 1/1000), anti-SOX9 (AB5535, Millipore, Burlington, MA, USA, 1/1000), anti-phospho-ERK (9101S, Cell Signaling Technology, 1/1000) and anti-total ERK (4695, Cell Signaling Technology, 1/1000). After washing, membranes were incubated with anti-rabbit 800CW or anti-mouse 680RD secondary antibodies (Li-Cor Biosciences) for 1 h at room temperature. Detection of protein signal was performed using Li-Cor Odyssey Fc Imaging System.

### RNA sequencing and data analysis

75 bp single-end RNAseq of purified human dedifferentiated acinar and duct cells was performed on an Illumina NextSeq 500 platform at VIB, Nucleomics Core, Leuven Belgium with TruSeq library prep. Low quality ends (< Q20) were trimmed using FastX 0.0.13 (http://hannonlab.cshl.edu/fastx_toolkit/index.html). Reads shorter than 35bp after trimming were removed. Using FastX 0.0.13 and ShortRead 1.16.3 (Bioconductor), polyA reads (more than 90% of bases equal A), ambiguous reads (containing N), low quality reads (more than 50% of the bases < Q25) and artifacts reads (all but 3 bases in the read equal one base type) were removed. The preprocessed reads were aligned to the reference genome of Homo sapiens (GRCh3773) with Tophat v2.0.13 (25). Within- and between-sample normalization was corrected for using full quantile normalization with the *EDASeq* package from Bioconductor. The number of reads were counted in the alignment that overlapped with gene features using featureCounts 1.4.6 (26). FPKM values were determined by dividing for each sample the normalized counts by the total number of counts (in millions). Then for each gene the scaled counts were divided by the gene length (in kbp). As such we got the number of Fragments Per Kilobase of gene sequence and per Million fragments of library size. Data will be publicly available on EBI (accession number E-MTAB-9386). 100 bp paired-end RNA sequencing of *Mecom* KD versus wild-type 266-6 cells was performed using Kappa HMR Library prep at BrightCore, UZ Brussel, Brussels, Belgium on an Illumina Novaseq 6000. The read quality of the raw data was evaluated using FASTQC (http://www.bioinformatics.babraham.ac.uk/projects/fastqc/), which flags any potential abnormalities that may have occurred during library preparation or sequencing. The raw reads were mapped against the Mus Musculus genome (version GRCm38-83) using the RNA-seq aligner STAR (27). The mapped reads were then translated into gene counts with the open-source tool (HTSeq) (28). Following quantification of expression levels, we carried out a differential expression analysis for both RNAseq datasets between experimental groups using the DESeq2 package in R (29). KEGG pathway analysis was performed using KOBAS gene set enrichment tool. Transcription factors were annotated using the Animal transcription factor database of human transcription factors.

### Biospecimens and clinical data

Biospecimens, transcriptomic and clinical data were collected by the Australian Pancreatic Cancer Genome Initiative (APGI), and informed consent was obtained from all patients. Use of clinical samples and data were in accordance with national ethical guidelines and regulations in Australia (HREC/11/RPAH/329—Sydney Local Health District—RPA Zone, protocol X11-0220) and in Belgium (UZ Brussel, B.U.N.143201732468).

### Statistics

Sample sizes were determined based on the type of sample, being 3 independent biological repeats for cell lines and 3-6 independent biological repeats for mice experiments. Specifically for the *in vivo* experiments with ElaCreERT and ElaCreERT;Mecom^f/f^ mice, sample size was calculated based on power analysis with power of 0.8 at a p=0.05 using G*Power (30). Statistical analysis was performed using Prism v8.0 (GraphPad Software, La Jolla, CA, USA). Differences among groups were tested for statistical significance by applying paired or unpaired two-tailed parametric Student’s t-test, or One-Way or Two-Way ANOVA with Bonferroni post-hoc test to correct for multiple comparisons. For each experiment, the specific statistical test is mentioned in the Figure legend. The data met the assumptions of the test and the variance between groups that were being statistically compared was similar. The number of independent repeats is indicated as N. Results are presented as mean ± SD. *p<0.05, **p<0.01, ***p<0.001 and ****p<0.0001.

## Results

### MECOM is a cancer-related transcription factor uniquely expressed in human dedifferentiated acinar cells

Given the difference in propensity of dedifferentiated acinar cells and duct cells for pancreatitis-associated tumor development (5), we were specifically interested in the differential gene expression of these two populations. A mixed human acinar and duct cell fraction was obtained as residual material from a pancreatic islet donor program. The cells were fluorescently labeled with the acinar-specific FITC-UEA-1 lectin and cultured in suspension, as described previously (12). After culture we FACS-purified dedifferentiated acinar cells and duct cells, based on FITC-UEA-1 and the duct cell marker CA19.9, respectively (11, 12) (Figure 1a). We confirmed the mutually exclusive expression pattern of CD142, a progenitor marker (11) in the UEA1+ dedifferentiated acinar cells, and of CA19.9 in the UEA1-duct cells (Figure 1b). We validated the different expression levels of the acinar cell markers MIST-1 and chymotrypsin, the progenitor marker CD142 and the duct-cell marker CK19 in the original mixed exocrine cell fraction and in the purified dedifferentiated acinar cells (UEA1+/CA19.9-) and native duct cells (UEA1-/CA19.9+) (Figure 1c). Comparing dedifferentiated acinar cells with native duct cells by RNAseq analysis highlighted 7953 differentially expressed genes (Adj. p<0.01). Of these, 1219 genes were expressed higher in dedifferentiated acinar cells (log fold change (FC) of ≤-2; Adj. p<0.01) (Supplementary Table 1) and were enriched for the KEGG pathways (Adj. p<0.01): ‘Protein digestion and absorption’ and ‘Pancreatic secretion’. This list included genes such as carboxypeptidase A and amylase, pointing to gene expression reminiscent of the abundant acinar cell transcriptome (31). In addition, ‘Pathways in cancer’ belonged to the top enriched pathways such as those of the WNT signaling pathway, which is well-known for its function in pancreatic tumorigenesis. Other enriched pathways included ‘PI3K-AKT signaling’, which is involved in acinar cell dedifferentiation and pancreatic tumorigenesis (32–34) and ‘Complement and coagulation cascades’ encompassing genes belonging to the tissue damage pathway (Figure 1e) and that code for F8, fibrinogen and F3 (CD142), the marker used above (11) (Figure 1b).

**Figure 1.**
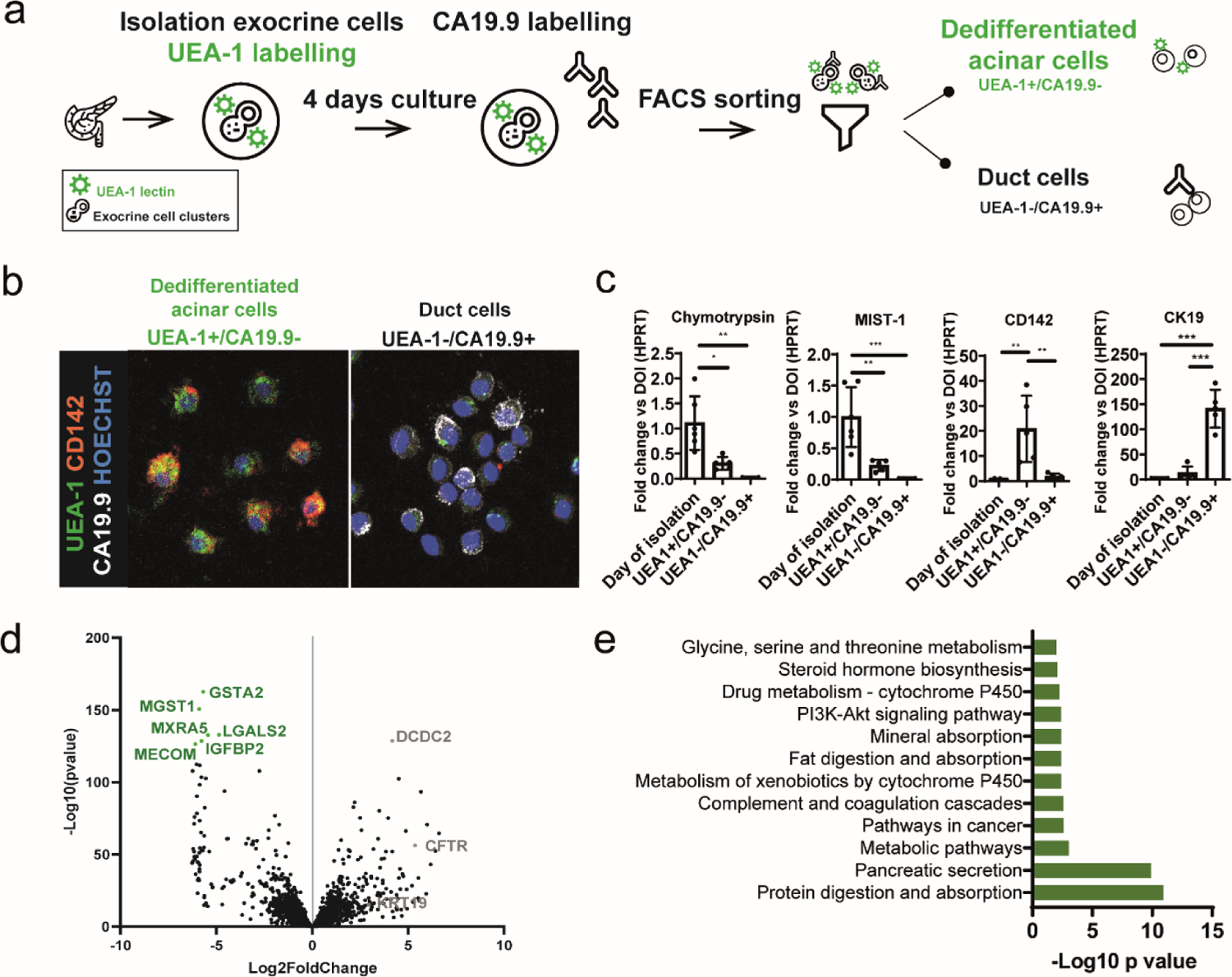
RNAseq analysis identifies *MECOM* as uniquely expressed in dedifferentiated acinar cells. (a) Schematic overview of FACS-purification of human dedifferentiated acinar and duct cells, using acinar-specific UEA-1 lectin. (b) Immunofluorescent staining of FACS-purified dedifferentiated acinar (UEA-1+/CA19.9-) and duct (UEA-1-/CA19.9+) cells at day 4 with CD142 and CA19.9. DNA is labelled in blue. (c) qPCR validation of RNAseq data of purified human dedifferentiated acinar and duct cells: Fold change expression (with HPRT as housekeeping gene) of purified dedifferentiated acinar cells (UEA-1+/CA19.9-) and duct cells (UEA-1-/CA19.9+) after day 4 of culture compared to the mixed exocrine fraction on day of isolation (day 0) (Mean ± SD; N = 5; *p<0.05; **p<0.01; ***p<0.001; one-way ANOVA with post-hoc Bonferroni correction). (d) Volcano plot of differentially expressed genes in human dedifferentiated acinar cells (green) and duct cells (grey). (e) KEGG pathways enriched (Adj. p-value < 0.01) in human dedifferentiated acinar cells (N = 5).

Dedifferentiated acinar and duct cells did have equal expression of the embryonic precursor gene *PDX1*, confirming our previous work (13). *PTF1A*, *NR5A2*, *RBPJL*, *FOXA3* and *GATA4* were expressed higher in the dedifferentiated acini, reminiscent of their fully differentiated state. Of note, *GATA6* expression, which is required for embryonic acinar cell differentiation (35, 36) and is a suppressor of pancreatic carcinogenesis (37, 38), was not different from the duct cells.

Conversely, duct cell-enriched genes included *CFTR*, *SOX9*, *NKX6.1*, *HES1*, *HNF1B* and *ONECUT2* (*HNF6B*), established markers of duct cell differentiation. The most significantly duct cell-enriched gene was *DCDC2* (Doublecortin Domain Containing 2), demonstrated to bind tubulin and enhance microtubule polymerization (39) (Figure 1d). In the duct cell fraction, we found KEGG pathway enrichment for ‘ECM receptor pathway interaction’ and ‘O-linked glycosylation’. Also, ‘Pathways in cancer’ featured here, but the genes in this pathway differed from the ones in the dedifferentiated acinar cell fraction (Supplementary Table 1).

There were 71 genes encoding transcription factors within the differentially expressed genes. The most significantly enriched transcription factor in the dedifferentiated acinar cell fraction was *MECOM* (log2FC = −6.09; Adj. p = 1.04E-123) (Figure 1d, Supplementary Table 1), a known oncogene (40–42). It has been described as an activator of KRAS and glypican-1 in pancreatic cancer (20, 21) and, very recently, it was reported in relation to acinar cell-derived tumorigenesis (43). Hence, we confirmed *MECOM* expression in human PDAC samples (N = 3) (Supplementary Figure 1a). We also used RNAseq analysis of stage 1 and 2 PDAC (N=103) to differentiate low and high MECOM expressors. MECOM expression correlated positively with disease-free survival (Supplementary figure 1b). In the same cohort, we previously reported a similar association for SOX9 and showed that SOX9 associated with the classical subtype, a molecular subset of PDAC that has better prognosis. Interestingly, in that same study we had found a correlation in expression between *SOX9* and *MECOM* (Pearson R=0,49) (23). When now we performed pathway analysis of genes correlating with MECOM (Supplementary figure 1c), indeed, similar pathways as those reported in our study on SOX9 were enriched, among which ErbB signaling was highlighted (23).

In conclusion, we FACS-purified human dedifferentiated acinar and duct cells and analyzed their transcriptome. The dedifferentiated acinar cell fraction showed a signature reminiscent of the acinar cell identity and specifically expressed *MECOM*, a transcription factor that featured in the pathway ‘Pathways of cancer’. MECOM was expressed in PDAC, where it correlated with SOX9 and ErbB/ERK signaling. We hypothesized that MECOM plays a role during acinar cell dedifferentiation, an event that initiates pancreatic tumor development, and this in connection to SOX9 and ERK signaling.

### MECOM is expressed in developing acinar cells and is re-expressed during pancreatitis-associated acinar cell dedifferentiation

We comprehensively assessed expression of MECOM during acinar cell differentiation or loss thereof. First, we confirmed the differential expression of *MECOM* in the purified human dedifferentiated acinar and duct cells from the experiments above (Figure 2a). *MECOM* mRNA appeared at low levels in the acinar compartment of normal human pancreas (0.24 ± 0.09 mRNA dots/cell; N=3), but increased in samples of chronic pancreatitis (1.46 ± 0.10 mRNA dots/cell; N=3; *p = 0.02) (Figure 2b,c).

**Figure 2.**
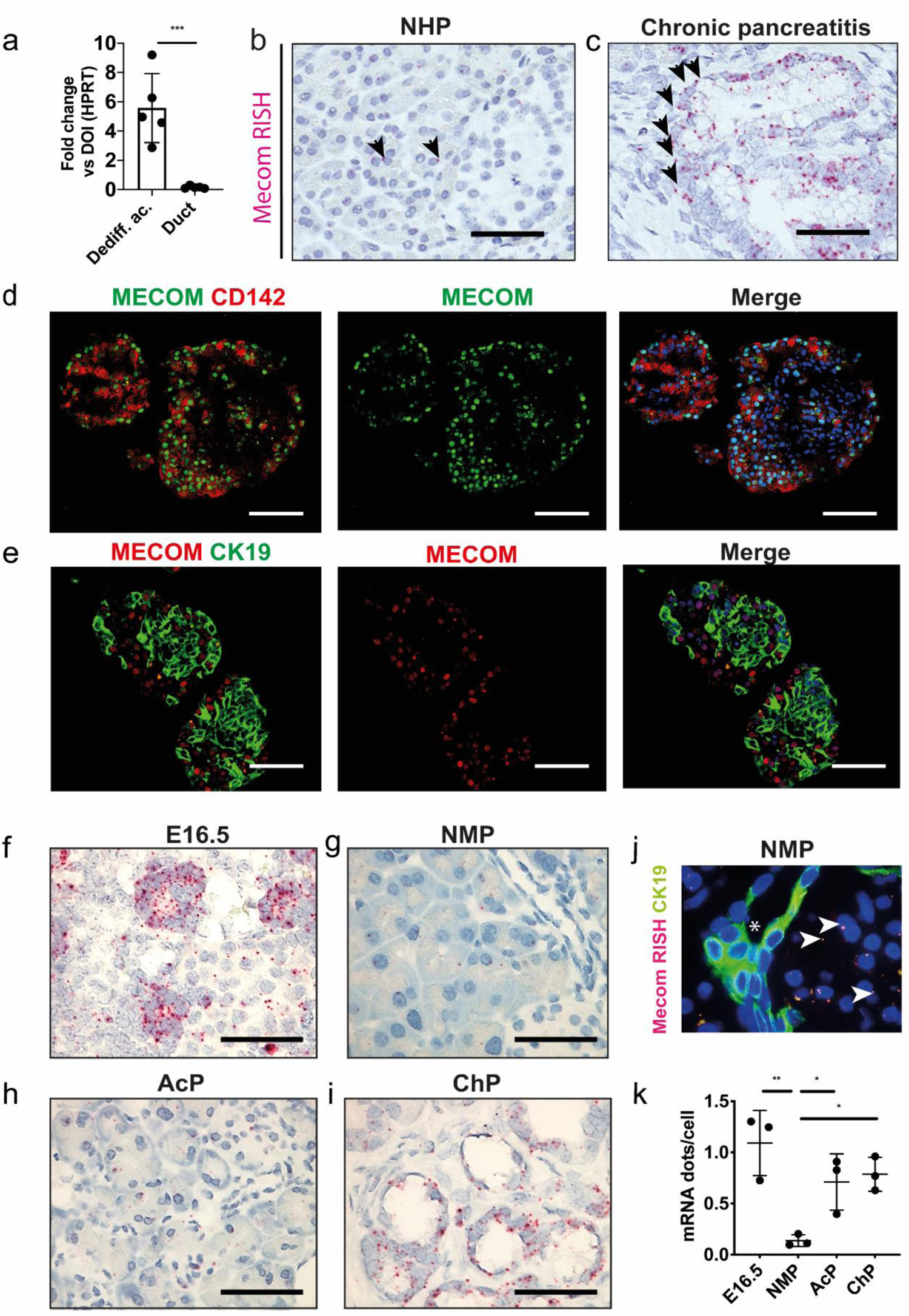
MECOM is uniquely upregulated in dedifferentiated acinar cells and not in duct cells. (a) qPCR for MECOM on purified human dedifferentiated acinar (UEA-1+/CA19.9-) versus duct cells (UEA-1-/CA19.9+) after FACS isolation at day 4 of culture, compared to the mixed exocrine fraction at day of isolation (day 0, D0) (N = 5; mean ± SD; ***p<0.001; One-way ANOVA with post-hoc Bonferroni correction). (b,c) RNA-ISH for MECOM on tissue from normal human pancreas (b) and human chronic pancreatitis (c) Scale = 50µM. Arrows indicate individual mRNA dots. (d,e) Immunofluorescent co-staining of human mixed exocrine fraction at day 4 of culture for MECOM and CD142 (d) and for MECOM and CK19 (e). Scale = 50µM. (f-i) RNA-ISH for *Mecom* and (k) expression level in normal mouse pancreas (NMP), E16.5 embryonic pancreas (g); and in a mouse model for acute (AcP) (h) and chronic (ChP) (i) pancreatitis. Scale = 50µM. (j) RNA-ISH for Mecom and immunofluorescent staining for CK19 in normal healthy mouse pancreas. The asterisk marks a pancreatic duct. (k) 5 individual microscopic fields per replicate were quantified (20x magnification) (N = 5; mean ± SD; *p<0.05; **p<0.01; One-way ANOVA with post-hoc Bonferroni correction).

Within the human purified dedifferentiated acinar cell fraction, *MECOM* showed strong correlation with *CD142* (Pearson R = 0.926, adj.P < 0.01), as discussed before (Figure 1). Immunofluorescent co-staining of MECOM and CD142 after 4 days of culture showed strong colocalization of the two proteins (56.94 ± 10.63% MECOM+/CD142+ cells; N = 5), while MECOM did not co-localize with the duct cell marker CK19 (5.69 ± 2.32% MECOM+/CK19+ cells; N = 5) (Figure 2d,e).

Next, we assessed the expression pattern (Figure 2f-i) and levels (Figure 2k) of *Mecom* mRNA in murine pancreas. In contrast to adult mouse pancreas, where *Mecom* was expressed at very low levels, a significantly increased expression was noted in E16.5 pancreas where it localized to pro-acinar tip cells (44, 45). Alike the human duct cells, we did not detect *Mecom* in CK19+ mouse duct cells (Figure 2j). Also similar to human, *Mecom* became re-expressed during acute and chronic pancreatitis in mice (Figure 2h,i,k).

Altogether, these data show that *Mecom* is transiently expressed in developing acinar cells and becomes re-expressed during pancreatitis in dedifferentiated acinar cells.

### SOX9 regulates *Mecom* expression in dedifferentiated acinar cells

SOX9 is an important marker of acinar cell dedifferentiation and is indispensable for tumor formation in the pancreas (23). SOX9 correlates with MECOM in PDAC patients, as referred above, an observation also reported in other tissues (46, 47). Hence, we investigated whether these two transcription factors might correlate in the context of acinar cell dedifferentiation. Indeed, we observed a strong correlation between *MECOM* and *SOX9* (Pearson R = 0.83; *p = 0.015; N = 6) (Figure 3a,b) in the human exocrine cell cultures in which acinar cells gradually dedifferentiate, a finding which was confirmed at the protein level (Figure 3c).

**Figure 3.**
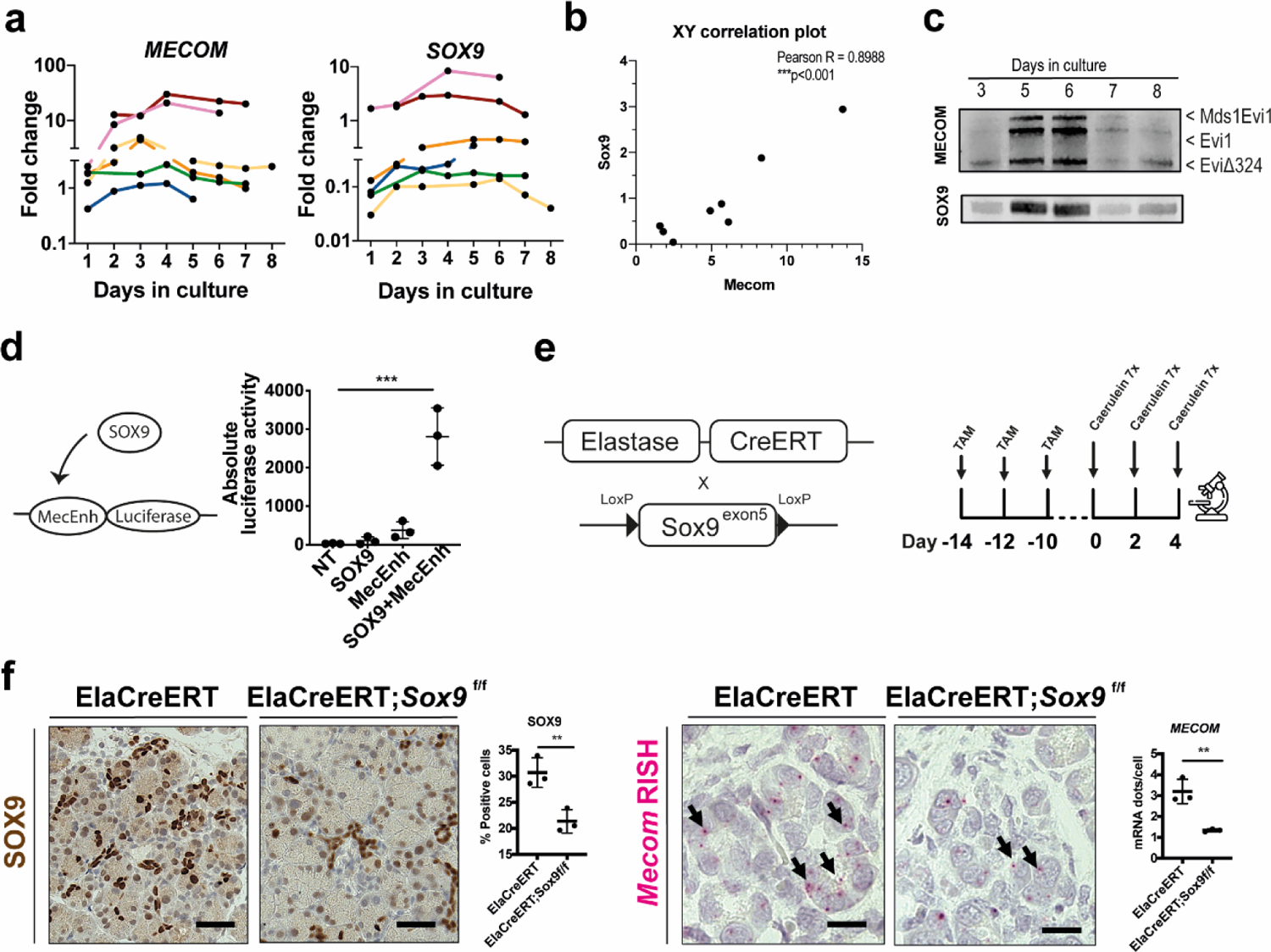
SOX9 regulates MECOM during acinar cell dedifferentiation. (a) qPCR and (b) XY correlation plot for MECOM and SOX9 expression on human exocrine mixed fraction in culture over time (day 1 to day 8). Each donor is indicated as a different colored line. (c) Immunoblot for MECOM and SOX9 on human exocrine mixed fraction in culture over time. Representative blot from N=3 human donor samples. The different isoforms of the MECOM locus are indicated by arrows. (d) Luciferase reporter assay in 266-6 cells, 48h after transfection. (mean ± SD; N = 3; ***p<0.001; one-way ANOVA with post-hoc Bonferroni correction). (e) Generation of acinar-specific (Elastase) Cre deletor mice with Sox9^ex5^ flanked by loxP sites. Acinar-specific deletion of SOX9 is performed in ElaCreERT;*Sox9*^f/f^ versus control (ElaCreERT) littermates by administering tamoxifen 3 times over a period of 5 days, followed by a two-week washout after start of tamoxifen treatment. Acute pancreatitis is induced by administering caerulein by 7 hourly injections for 3 days over a period of 5 days. Mice are sacrificed at day 4 after caerulein administration. Immunohistochemical staining and for SOX9 and RNA in situ hybridization of *Mecom* mRNA expression and quantification in mouse pancreatic tissue after experimentally induced acute pancreatitis in control (ElaCreERT) and acinar-specific Sox9 knockout (ElaCreERT;*Sox9^f^*^/f^) mice. Scale = 50µM. (Mean ± SD; N = 3; **p<0.01; unpaired two-tailed t-test)

To further investigate this mechanistically, we turned to the partially differentiated mouse acinar cell line 266-6 (that expresses *Mecom*, Figure 4a) where we performed a promoter-reporter assay. We observed low increase in luciferase activity when cells were transfected with the *Mecom* enhancer expression plasmid alone (MecEnh). After combined *Sox9* overexpression, luciferase activity increased significantly (SOX9 + MecEnh) (Figure 3d), indicating that Sox9 regulates the *Mecom* enhancer in partially differentiated acinar cells. Based on the observed induction of *Mecom* after 48 hours of caerulein-induced acute pancreatitis (Figure 2h, k), we investigated *Mecom* expression in *Sox9* loss-of-function mice under similar conditions (Figure 3e, f) (13, 48). ElaCreERT;*Sox9*^f/f^ mice showed 3.20 ± 0.58 Mecom mRNAdots/cell versus 1.34 ± 0.07 in control mice (ElaCreERT) (58.14 ± 8.46% reduction, **p = 0.0054) (Figure 3f), re-affirming that SOX9 acts as an upstream regulator of *Mecom*. In conclusion, we showed that *Mecom* is regulated by SOX9 in pancreatic acinar cell context.

**Figure 4.**
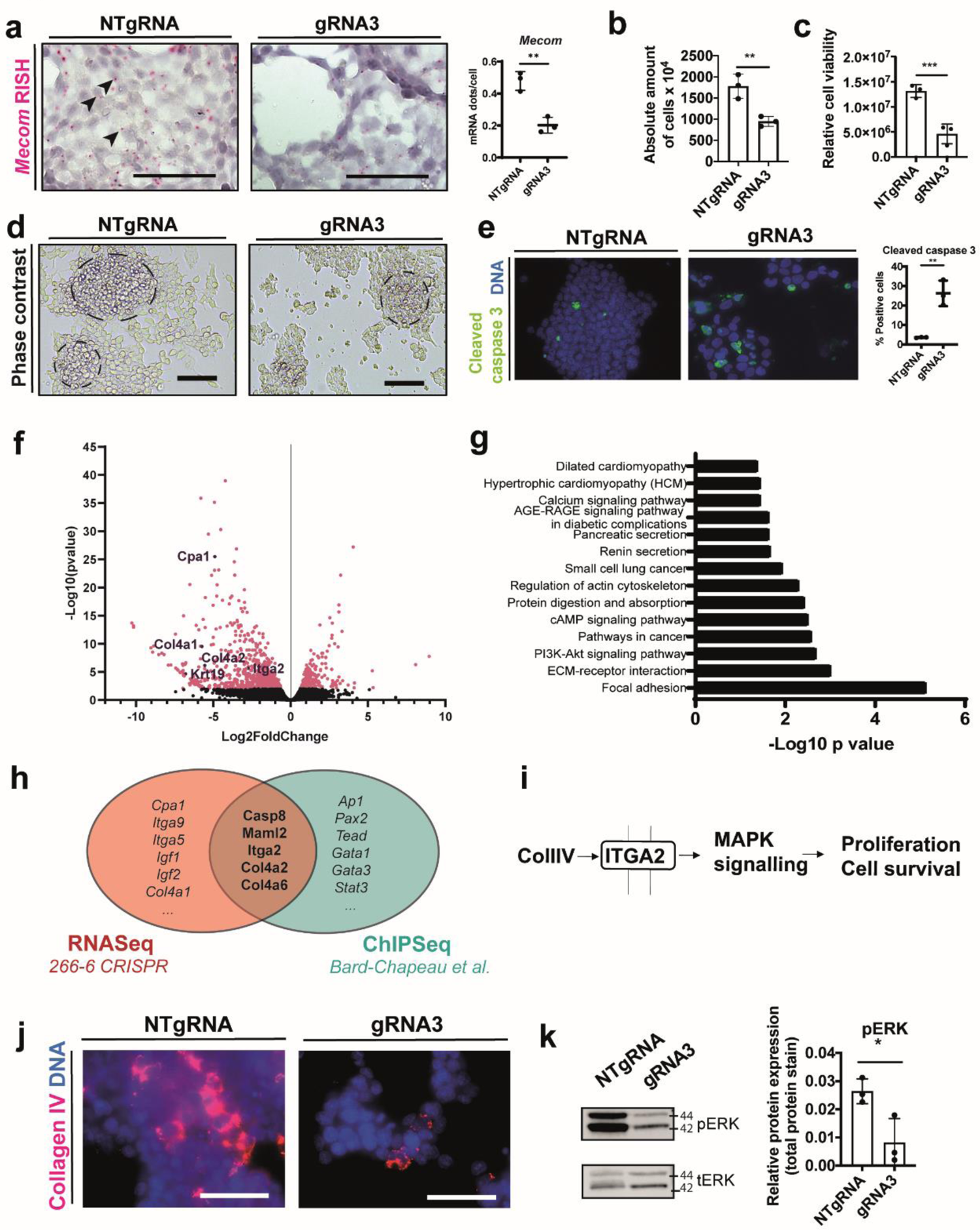
*Mecom* retains cell adhesion and survival in 266-6 cells. (a) RNA in situ hybridization (RISH) and quantification on control (NT-gRNA) and *Mecom* KD (gRNA3) 266-6 cells. Scale = 200µM. (N = 3; **p<0.01; unpaired two-tailed t-test). (b) Phase contrast images of control (NT-gRNA) and *Mecom* KD (gRNA3) 266-6 cells. Scale = 200µM. (c) Absolute number of cells at 96 h after seeding 50 000 cells (mean ± SD; N = 3; **p<0.01; unpaired two-tailed t-test) (d) Cell viability measured relative to background levels of the culture medium by CellTiter Glo assay (mean ± SD; N = 3; ***p<0.001; unpaired two-tailed t-test) (e) Immunofluorescent staining and quantification of cleaved caspase 3+ cells. (N = 3; **p<0.01; unpaired two-tailed t-test) (f) Volcano plot and (g) KEGG pathway analysis on differential gene expression profiles after RNAseq analysis of *Mecom* wildtype versus *Mecom* KD cells (N = 3) (h) Venn diagrams showing genes overlapping between differentially expressed genes in *Mecom* KD 266-6 cells and publicly available ChIP-seq dataset from Bard-Chapeau et al. (49) (i) Schematic representation of our hypothesis. (j) Collagen IV immunofluorescent staining in red on 266-6 NTgRNA and gRNA3-transduced cultured cells. (k) Immunoblot for phospho-ERK and total ERK and quantifications of pERK relative to total protein stain. (mean ± SD; N = 3; *p<0.05; Unpaired two-tailed t-test)

### *Mecom* maintains cell adhesion and cell viability in a partially differentiated acinar cell line and in human and mouse acinar cell cultures

We engaged in a lentiviral all-in-one CRISPR/Cas9-mediated approach with 3 different gRNAs designed to knock down *Mecom* in a partially differentiated acinar cell line (266–6) to study the role of *Mecom* in acinar cell dedifferentiation. Only gRNA3 provided evaluable knock-down (KD) (Supplementary Figure 2) and was used further. RISH for *Mecom* in this loss-of-function condition (Figure 4a) confirmed significant reduction of mRNA expression (0.48 ± 0.06 *Mecom* mRNA dots/cell in *Mecom* KD cells compared to 0.20 ± 0.05 *Mecom* mRNA dots/cell in control cells; 55.50 ± 14.43 % reduction; **p = 0.0033). In the *Mecom* KD condition, the cells were not able to organize into the typical ‘acinar dome-like’ structures and seemed to lack the capacity to attach properly to the culture dish compared to the cells transduced with non-targeting gRNA (NTgRNA) (Figure 4b), suggestive of loss of proper cell-cell contacts. We also observed a decrease in absolute cell number (Figure 4c), an overall diminished cell viability (Figure 4d) and a significant increase in cleaved caspase 3+ cells (Figure 4e), all indicative of increased cell death and loss of adhesive capacity.

We further studied effects of *Mecom* KD in this cell line by RNAseq analysis. Differential gene expression analysis (Figure 4f) revealed downregulation of genes enriched in ‘Focal adhesion’, ‘ECM receptor interaction’ and ‘Pathways in cancer’ (Figure 4g), underscoring our observations of diminished dome formation. We also compared the differentially expressed genes from the *Mecom* KD condition with a publicly available MECOM ChIP-Seq dataset (49) and found 9.89% overlapping genes (Figure 4h) to make an informed decision on which genes to consider further. In particular, we assessed integrins that link the extracellular matrix (ECM) components such as collagen to the cytoskeleton, hypothesizing that *Mecom* might compromise cell-cell interaction by diminished Collagen IV-Integrin a2 interaction and downstream ERK-mediated signaling (Figure 4i), the latter considered crucial in acinar cell dedifferentiation (50). Diminished expression of collagen type IV and reduced ERK signaling upon *Mecom* loss-of-function was confirmed by immunofluorescent staining (Figure 4j,k).

In conclusion, *Mecom* is indispensable to maintain cell adhesion and cell viability in a partially differentiated acinar cell line. Next, we crossed ElaCreERT deletor mice with mice carrying a floxed exon 4 of *Evi*, an exon which is conserved in all transcript variants (24) (Figure 5a). We studied effects in acinar cells isolated from these mice using an established cell culture model of acinar cell dedifferentiation, same as the human cell culture model described above (13). Immunohistochemical staining for MECOM showed 42.71 ± 13.42% less MECOM positive cells in freshly isolated (day 0) MECOM KO exocrine cells, consistent with previous reports on the recombination efficiency of the ElaCreERT strain (51–53). Interestingly, by day 4, we detected only 23.89 ± 13.42% less MECOM positive cells in the KO condition suggesting selective loss of MECOM deficient cells (Supplementary Figure 3).

**Figure 5.**
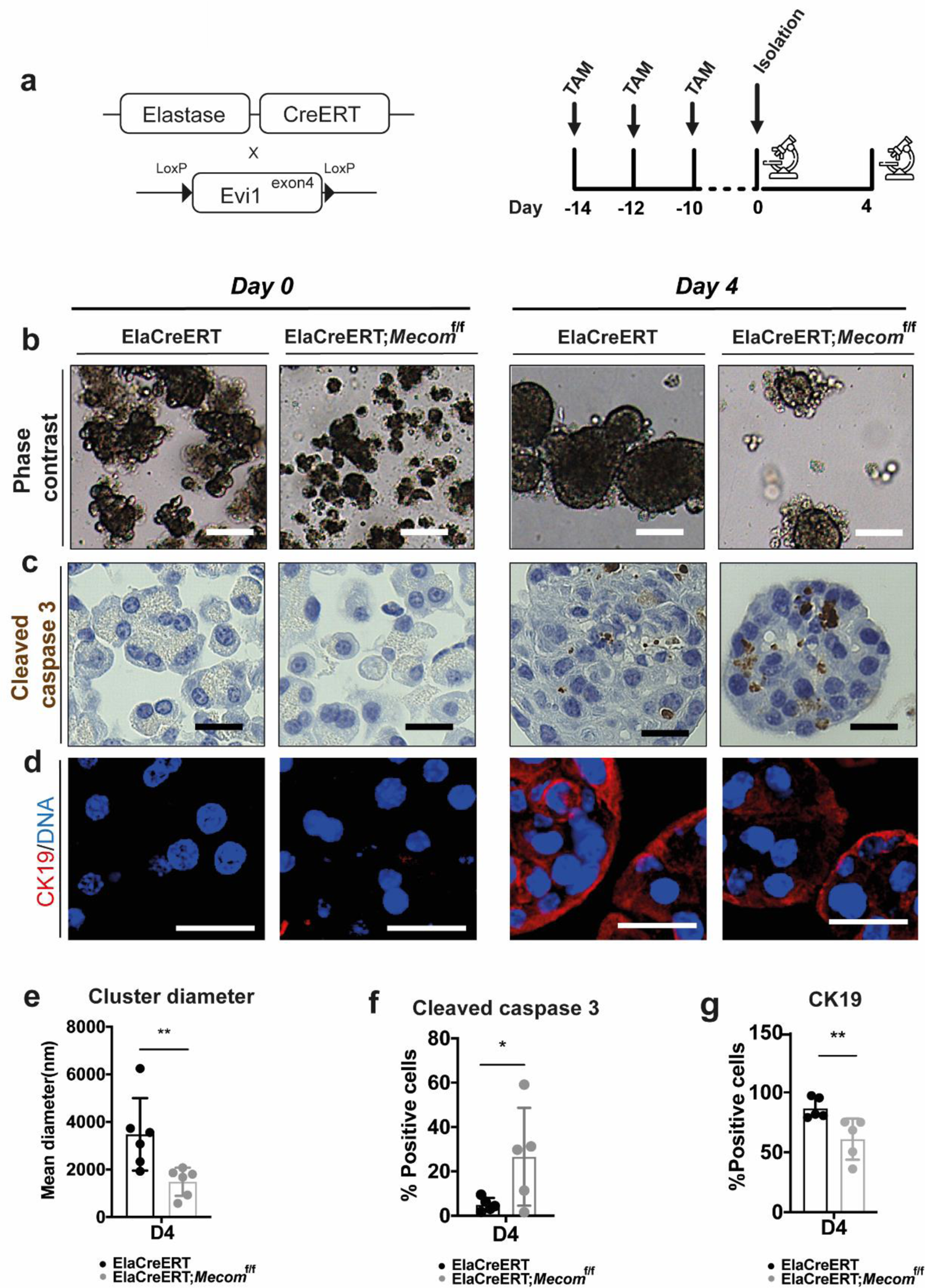
*Mecom* retains cell adhesion and survival in cultured acinar cells. (a) Acinar cell-specific deletion of MECOM is performed by administering tamoxifen 3 times over a period of 5 days both subcutaneously and by gavage, followed by a two-week washout period after the start of tamoxifen treatment. Acute pancreatitis is induced by administering caerulein by 7 hourly injections for 3 days over a period of 5 days. Mice are sacrificed at day 4 after caerulein treatment. (b,e) Phase contrast images and quantification of mean cluster surface of ElaCreERT and ElaCreERT;*Mecom*^f/f^ isolated acinar cells immediately after isolation (D0) and at day 4 of culture (D4). Scale = 100µM. (c,f) Cleaved caspase 3 staining and quantification on ElaCreERT and ElaCreERT;*Mecom*^f/f^ acinar cells at day 4 of culture. Scale = 25µM. (d,g) CK19 staining and quantification of ElaCreERT and ElaCreERT;*Mecom*^f/f^ acinar cells at day 4 of culture. Scale = 25µM. (mean ± SD; N = 6; *p<0.05; **p<0.01; two-way ANOVA with post-hoc Bonferroni correction.)

We observed that after cell isolation the control cells remained organized in small acinar spheroid-like structures while in the MECOM KO condition they often presented as single cells. After 4 days of culture, clusters did form in the ElaCreERT;*Mecom*^f/f^ mouse cells but cluster diameter was significantly smaller compared to those of ElaCreERT control mice (57.21 ± 15.73% reduction in average diameter) (Figure 5b,e), suggestive of diminished cell adhesion and cell death of the single cells. In our experience, isolated acinar cells cannot survive as single cells (e.g. after FACS purification).

Similar to our results in the 266-6 cells (Figure 4), an increased fraction of acinar cells in the MECOM KO cultures stained positive for cleaved caspase 3 (81.21 ± 26.47% increase in cleaved caspase 3+ cells) (Figure 5c,f). Notably, the MECOM KO cultures also showed less CK19+ cells in comparison to the control cells (25.79 ± 6.33% reduction in CK19+ cells), suggesting a lower capacity for dedifferentiation (Figure 5d,g).

These observations were confirmed by shRNA-mediated knockdown in cultured human exocrine cells in culture. At 24h, 4 and 7 days after transduction, average cluster diameter was significantly decreased (Supplementary Figure 4), underscoring the compromised cell-cell adhesion.

In conclusion, MECOM is indispensable for maintenance of cell-cell adhesion and cell viability in primary cultures of mouse and human acinar cells.

### Mice with *Mecom*-deficient acinar cells show more acinar cell death and prolonged immune infiltration after acute pancreatitis

Since the primary cell culture models above recapitulate acinar cell dedifferentiation as it occurs during pancreatitis (11–13), we subjected control and MECOM KO mice to caerulein-induced acute pancreatitis. In wild-type mice, acute pancreatitis peaks around day 4 and resolves by day 11 (Figure 6a). We observed no macroscopic differences in the pancreas nor in weight of the pancreas or body weight when comparing MECOM KO with control mice (Supplementary Figure 5). Haematoxylin-eosin staining however showed more inter-acinar spaces at day 11 in the MECOM KO mice (Figure 6b, f). Similar to the phenotype observed in the 266-6 cells (Figure 4) and the *ex vivo* cultures (Figure 5), this was preceded by an increased incidence of apoptosis (Figure 6c,g). Since we observed more inter-acinar space in the MECOM KO mice, we analyzed immune cell infiltration. At day 11, CD3 positive area in MECOM KO mice remained at a level comparable to the acute phase of pancreatitis (day 4), whereas in the control mice it was significantly decreased (Figure 6e,i). F4/80+ macrophages showed a similar trend, and CD19+ B-cells were only present in the surrounding lymph nodes (Supplementary Figure 6). Remarkably, no difference in Masson’s Trichrome blue staining could be detected between control and MECOM KO mice (Figure 6d,h), indicating that the inter-acinar space was not filled with collagen. The histological pattern showing accumulation of excessive liquid in the interstitial space was consistent with that of edema. Also, we did not observe any statistically significant changes in acinar-to-ductal transdifferentiation markers at day 11 (not shown).

**Figure 6.**
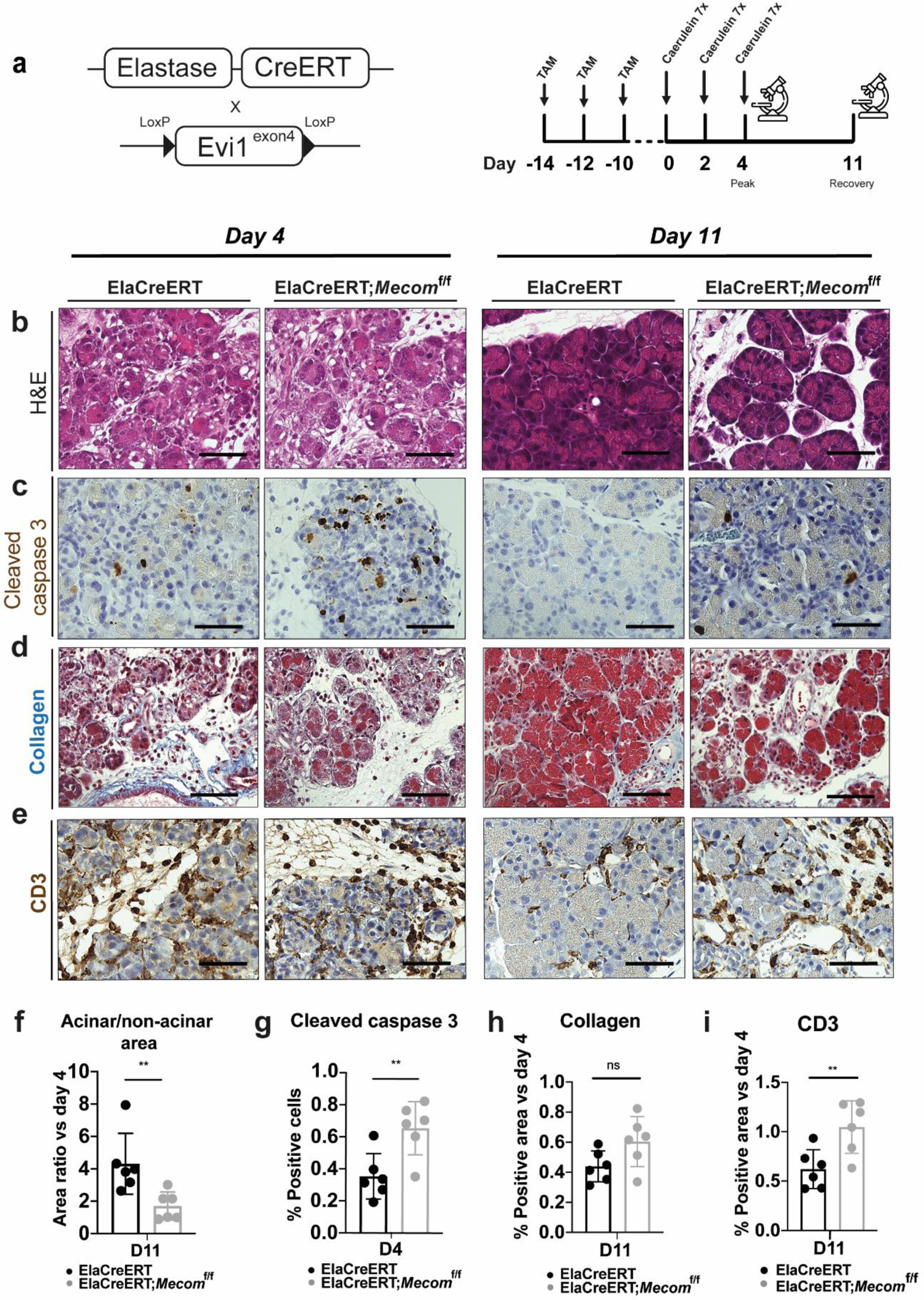
*Mecom* depletion in acinar cells *in vivo* causes pronounced cell death during recovery after caerulein-induced acute pancreatitis. (a) Acinar-specific deletion of MECOM is performed by administering tamoxifen 3 times over a period of 5 days, followed by a two-week washout period after the start of tamoxifen treatment. Acute pancreatitis is induced by administering caerulein by 7 hourly injections for 3 days over a period of 5 days. Mice are sacrificed at day 4 and day 11 after caerulein administration. (b,f) H&E staining and quantification of acinar/non-acinar area ratio relative to day 4. (c,g) Cleaved caspase 3 immunohistochemical staining and quantification at day 4. (d,h) Collagen staining and quantification relative to day 4. Scale = 50µM. (e,i) CD3 staining and quantification relative to day 4 for ElaCreERT and ElaCreERT;*Mecom*^f/f^ mice. Scale = 50µM. (mean ± SD; N = 6; **p<0.01; ns = non-significant. two-way ANOVA with post-hoc Bonferroni correction)

Altogether, these results indicate that MECOM-deficiency in pancreatic acinar cells under the experimental stress of acute pancreatitis leads to increased acinar cell death with ensuing immune cell infiltration and edema.

## Discussion

When subjected to stress, such as occurring during chronic pancreatitis, acinar cells dedifferentiate and acquire a ductal cell-like phenotype, rendering them prone to *Kras* induced transformation, in contrast to the native duct cells which seem more resistant to neoplastic development. Where studies in the past focused on the acinar cell plasticity and on how the dedifferentiated acinar cells become duct-like (13), the inherent differences between the two cell populations that may underly a different propensity to tumor formation remained unclear. Therefore, we profiled, for the first time to our knowledge, the transcriptome of human dedifferentiated acinar cells versus their native duct cell counterpart. This comparative transcriptomic analysis provided several new insights, including the fact that both cell populations were enriched in ‘Pathways of cancer’ although the related genes differed in both fractions. We identified MECOM, a transcription factor known to be involved in (pancreatic) cancer, as the most uniquely expressed gene in the dedifferentiated acinar cells.

MECOM is known to regulate many aspects of tumorigenesis including differentiation, proliferation and apoptosis (17–19, 46). It is a regulator of stomach-specific genes, some of which become upregulated in dedifferentiated acinar cells (23), and was identified in a specific cluster of acinar cells during PDAC development in a scRNAseq experiment recently published (54). Previously, a role for MECOM in pancreas cancer was proposed (20, 21, 43). Here, we focused on its role in acinar cell dedifferentiation. We uncovered that MECOM is uniquely expressed in dedifferentiated mouse and human acinar cells and absent from native duct cells, both in cell cultures and in tissue samples. Notably, MECOM expression levels appeared even higher in embryonic mouse pancreas, specifically in the pro-acinar tip cells again suggesting a relation to an immature acinar cell phenotype.

Interestingly, SOX9 is a known regulator of acinar cell dedifferentiation and a driver of pancreatic tumor formation (21, 55). Here, SOX9 expression correlated with that of MECOM. An interplay of MECOM and SOX9 has been described in other tissues (12, 13, 50) and we confirmed that SOX9 transactivates the *Mecom* enhancer in an acinar cell context. Still, it requires further investigation why SOX9 does not transactivate MECOM expression in duct cells where SOX9 expression is the highest. SOX9 likely needs other binding partners, that are absent in duct cells, to activate MECOM expression.

We observed a clear phenotype when culturing MECOM deficient acinar cells (266-6 cells and primary mouse and human acinar cells), i.e. impaired formation of acinar domes or cell clusters, suggesting impaired cell adhesion and higher rates of apoptosis. We showed that MECOM depletion reduced the cell-cell interactions through *Itga2* and its ligands such as *Col4a2*. In our *Mecom* KD cells, the expression of several integrins (*Itga2, Itga4, Itga5 and Itga9*) was decreased. Importantly, available ChIP-Seq data of MECOM in human ovarian carcinoma cells also showed direct binding to the promotor regions of *Col4a2* and *Itga2* (49). We confirmed this in *Mecom* KD cells, where reduced activity of the ERK signaling pathway could be directly linked to reduced acinar cell dedifferentiation, a process for which this signaling pathway is critical (12, 50). In a caerulein-induced acute pancreatitis model in MECOM-deficient mice, we again make similar observations and demonstrated a prolonged immune cell infiltration, similar as reported recently by Ye et al. (43).

We propose that normal acinar cells have two options when they are subjected to stress, either to subside in an “incognito” reversible dedifferentiated state, or to die. MECOM seems indispensable for the former option. Ye et al., who, like us, did not provide formal proof by lineage tracing, claimed that MECOM-deficient acinar cells transdifferentiate more extensively during pancreatitis (43), while we suggest that MECOM-deficient acinar cells undergo apoptosis and therefore cannot contribute any longer to the CK19+ transdifferentiated cell pool. This is supported by our observations increased inter-acinar spaces filled with immune infiltrates and fluid (edema) in the acute pancreatitis model. Such phenotype has also been reported in EGF knockout mice (56). We also did not detect any increase in collagen deposition around the recovering acini, which clearly differs from acinar-to-ductal transdifferentiation that is accompanied by typical collagen deposition (57), further supporting our conclusion.

Altogether, these results show that dedifferentiated acinar cells have a clearly distinct gene expression profile compared to native duct cells. Under stress, MECOM expression is turned on, downstream of SOX9, where it prevents acinar cell death and allowing cellular dedifferentiation.

## Acknowledgements

We thank Gunter Leuckx, Emmy De Blay, Veerle Laurysens, Geert Stangé, the laboratory of Liver Cell Biology Research Group at Vrije Universiteit Brussel, BrightCore at UZ Brussel and Flemish Institute of Biotechnology for expert technical assistance. We thank Patrick Jacquemin for providing the ElaCreERT and ElaCreERT;Sox9f/f mice, and Susumu Goyama and Mineo Kurokawa for providing the Evi-1 floxed mice.

## Author contributions

EB, EW, IH, LB and IR conceived and designed the research; EB, EW, JB, EM, MVB, YH and IH performed experiments; EB, EW, IH, LB and IR drafted the manuscript; DLDP, MK, SG and PJ contributed to animals and human donor material.

## Ethics approval

Biospecimens, transcriptomic and clinical data were collected by the Australian Pancreatic Cancer Genome Initiative (APGI, www.pancreaticcancer.net.au), which is supported by an Avner Pancreatic Cancer Foundation Grant (www.avnersfoundation.org.au). Informed consent was obtained from all patients. Use of clinical samples and data were in accordance with national ethical guidelines and regulations in Australia (HREC/11/RPAH/329—Sydney Local Health District—RPA Zone, protocol X11-0220) and in Belgium (UZ Brussel, B.U.N.143201732468). Consent for the use of residual donor material for research was obtained according to the legislation in the country of organ procurement. All animal experiments were approved by the Ethical Committee for Animal Testing at the Vrije Universiteit Brussel (#17-637-1).

