## Supplementary Figures for "MECOM permits pancreatic acinar cell dedifferentiation avoiding cell death under stress conditions"

**SUPPLEMENTARY DATA**

**Supplementary Table 1**. RNAseq and KEGG pathway analysis of differentially expressed genes in human dedifferentiated acinar cells and duct cells

**Supplementary Table 2**. RNAseq and KEGG pathway analysis of *Mecom* CRISPR KO in 266-6 cells

**Supplementary Figure 1. MECOM expression in human PDAC.** (a) RNA in situ hybridization for *MECOM* in human PDAC tissue (N=3). (b) Disease-free survival curve in PDAC patients with high and low MECOM expression (n=103). (c) Pathway analysis of genes correlated with MECOM in PDAC patients (N=103).

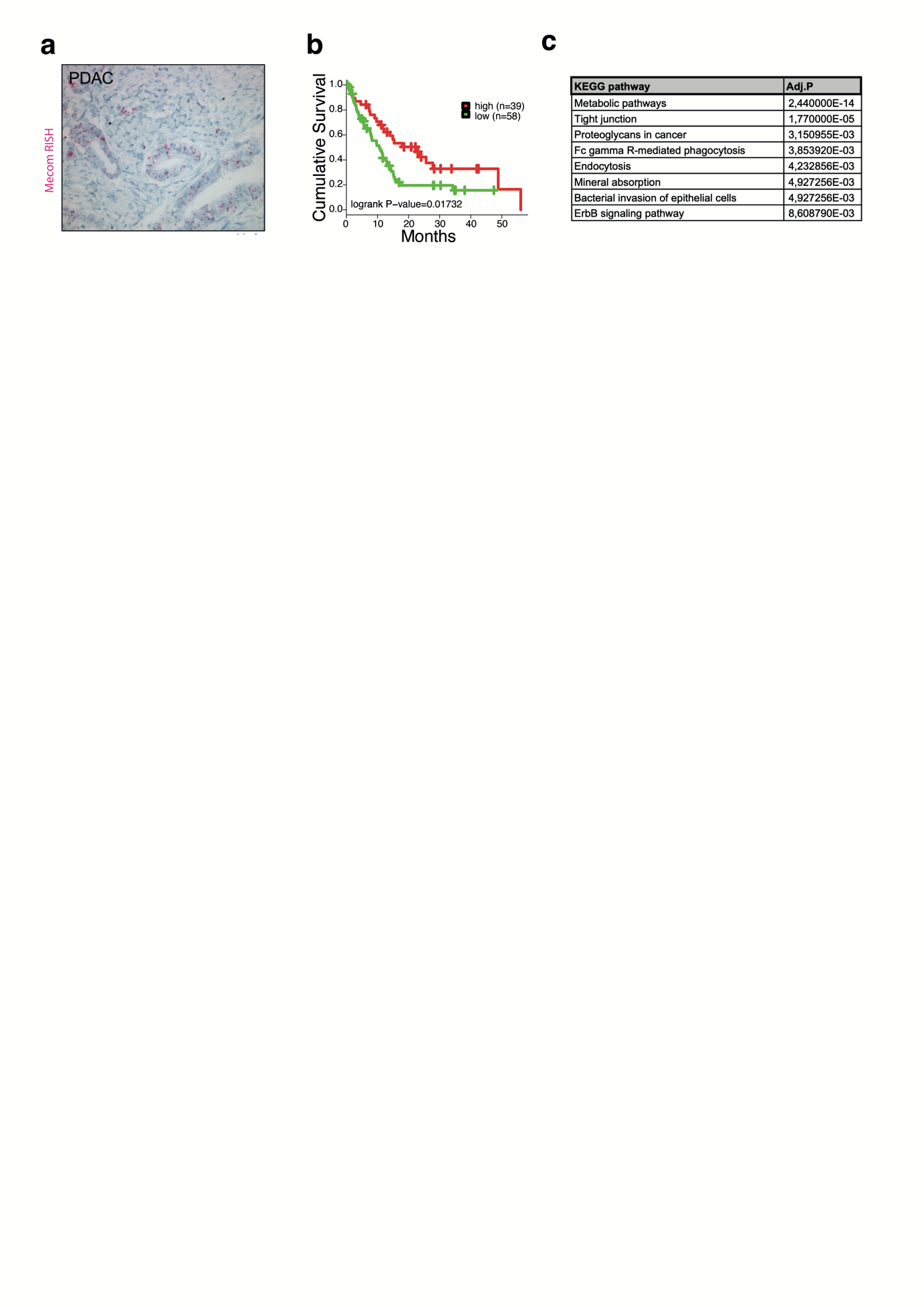

**Supplementary Figure 2.** CRISPR-mediated knockdown of *Mecom* in the partially differentiated mouse acinar cell line 266-6. Comparison of 3 different gRNAs designed to target mouse *Mecom* expression analyzed by qPCR. (mean ± SD; N=3)

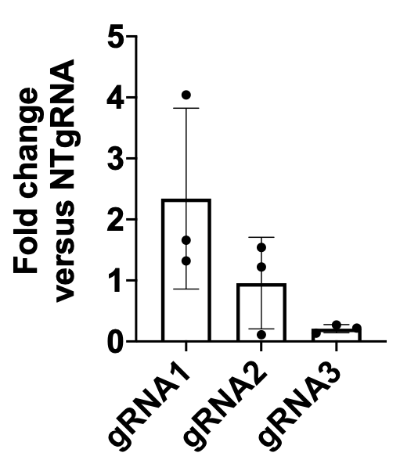

**Supplementary Figure 3.** Immunohistochemical staining and quantification of MECOM on ElaCreERT control and ElaCreERT;*Mecom*^f/f^ isolated acinar cells dedifferentiated in culture. Scale = 50M. (mean ± SD; N = 5; *p<0.05; two-way ANOVA with post-hoc Bonferroni correction)

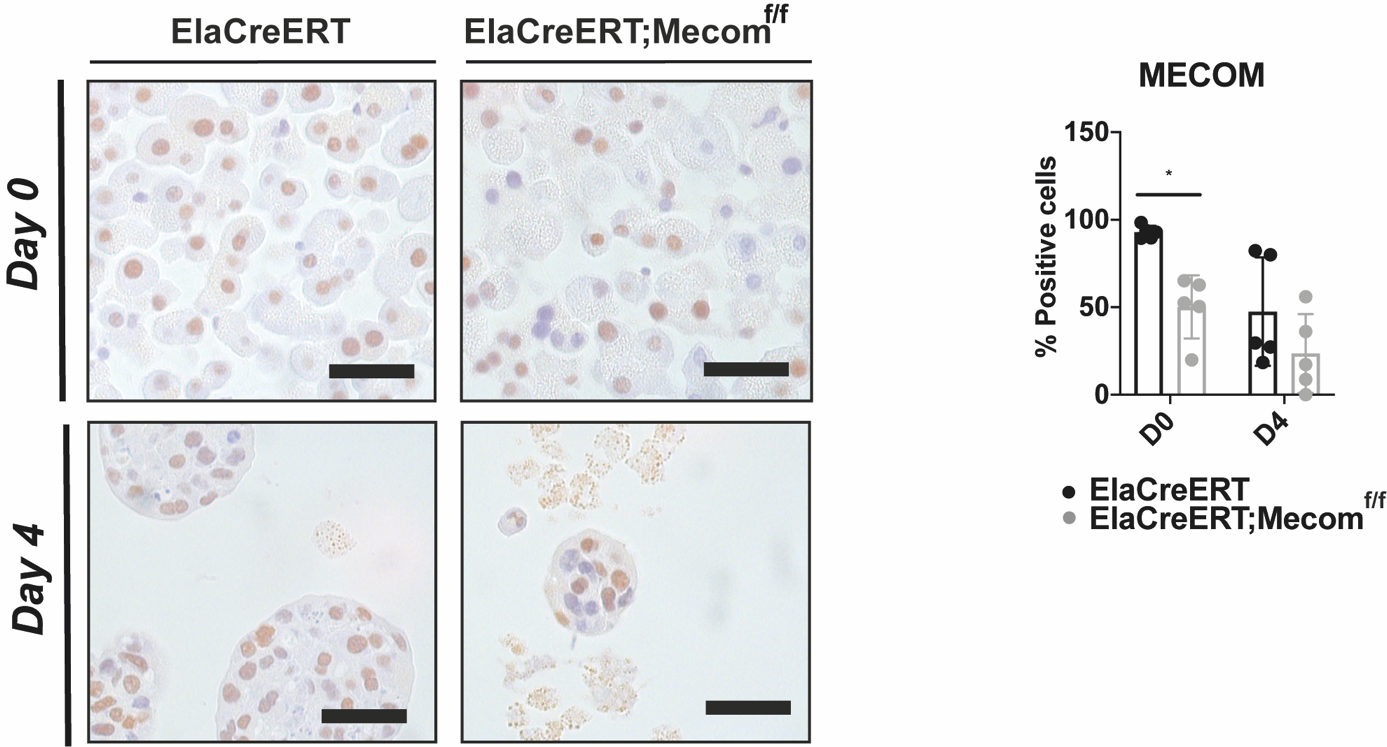

**Supplementary Figure 4. shRNA-mediated knockdown of *MECOM* in human exocrine cells dedifferentiating in culture:** (a) Representative images and cluster diameter of MECOM shRNA-transduced and non-transduced control human exocrine cells at 24h, 4 days and 7 days after transduction. Scale = 400µM. (mean ± SD; N = 3; ***p<0.001; ****p<0.0001; paired two-tailed t-test). Each datapoint represents a separate cluster. (b) qPCR of *MECOM* expression at day of isolation and 4 and 7 days after transduction in control and shRNA-transduced cultures. (mean ± SD; N = 3; *p<0.05; one-way ANOVA with post-hoc Bonferroni correction).

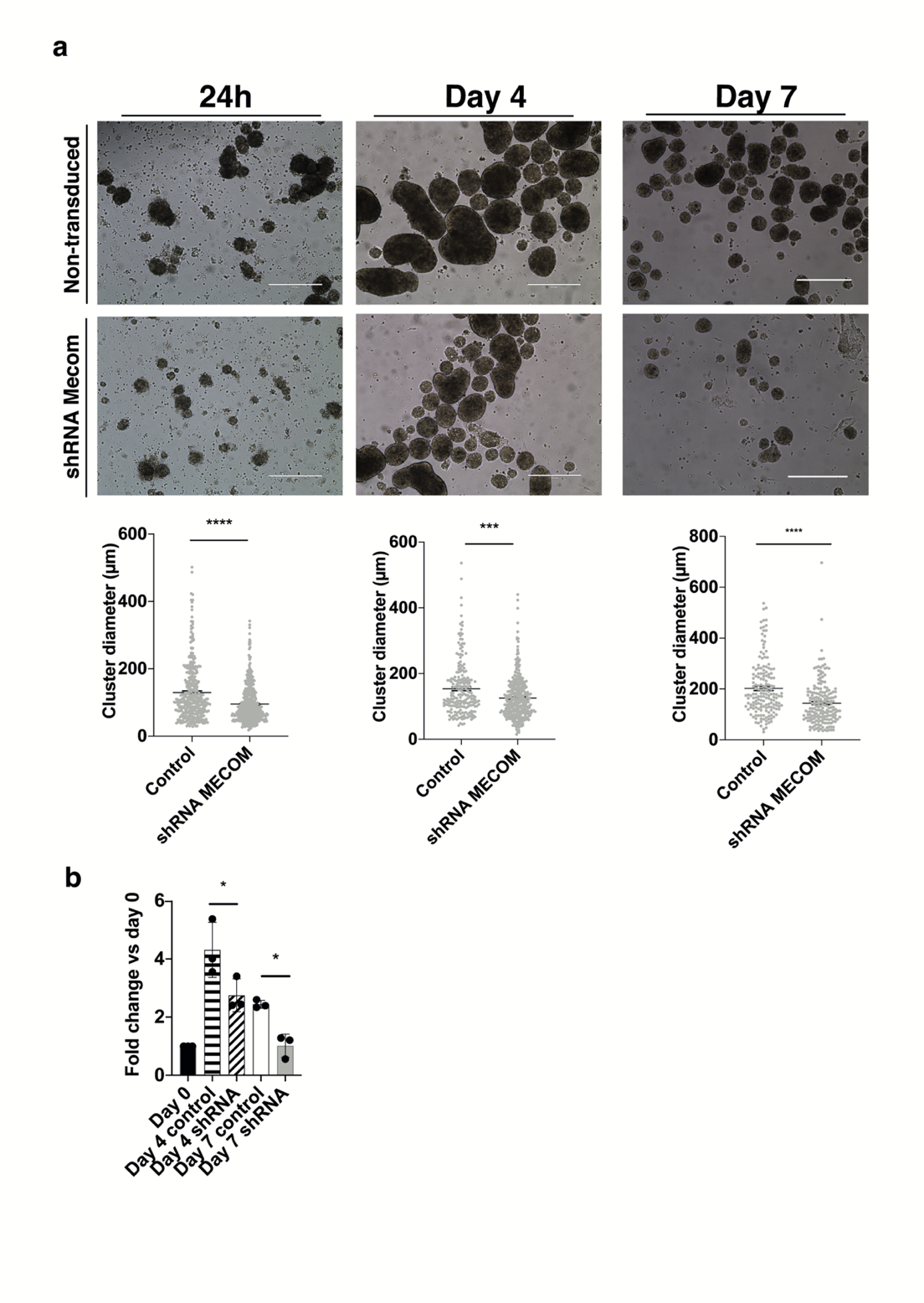

**Supplementary Figure 5**. (a) Pancreata, (b) body weight and (c) pancreas/body weight of control and ElaCreERT;*Mecom*^f/f^ mice at 4 (D4) and 11 days (Day 11) after the start of caerulein administration. (mean ± SD; N=6).

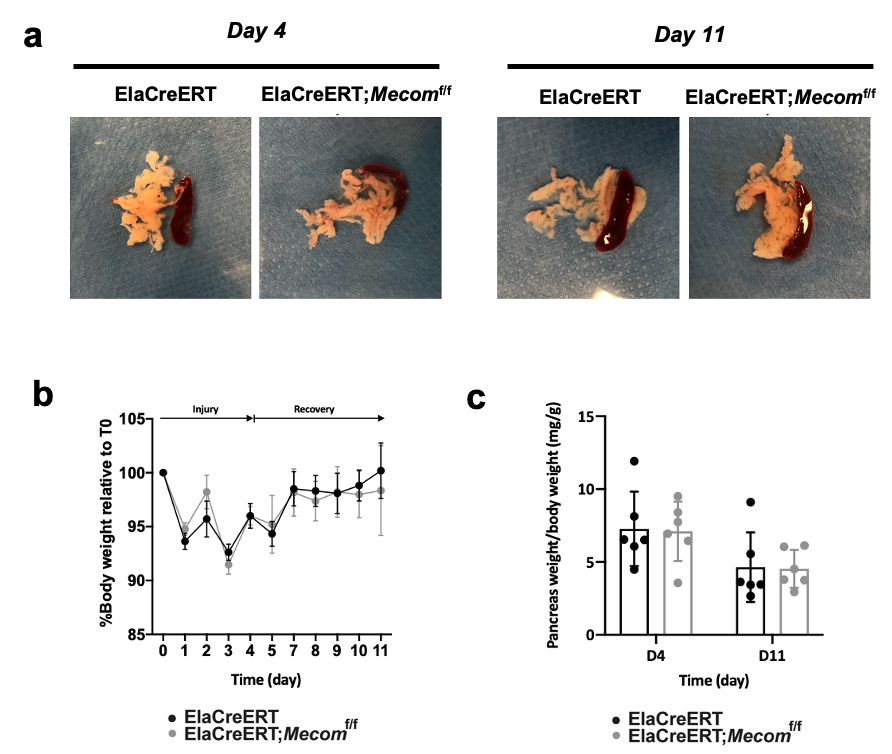

**Supplementary Figure 6**. **F4/80 and CD19 staining and quantification.** ElaCreERT and ElaCreERT;Mecom^f/f^ tissues sections after experimentally induced acute pancreatitis at the peak of inflammation (day 4) and after one week of recovery (day 11). Scale = 50µM. Mean ± SD; N = 6; ns = non-significant; two-way ANOVA with post-hoc Bonferroni correction). Lymph nodes are indicated with a white asterisk.

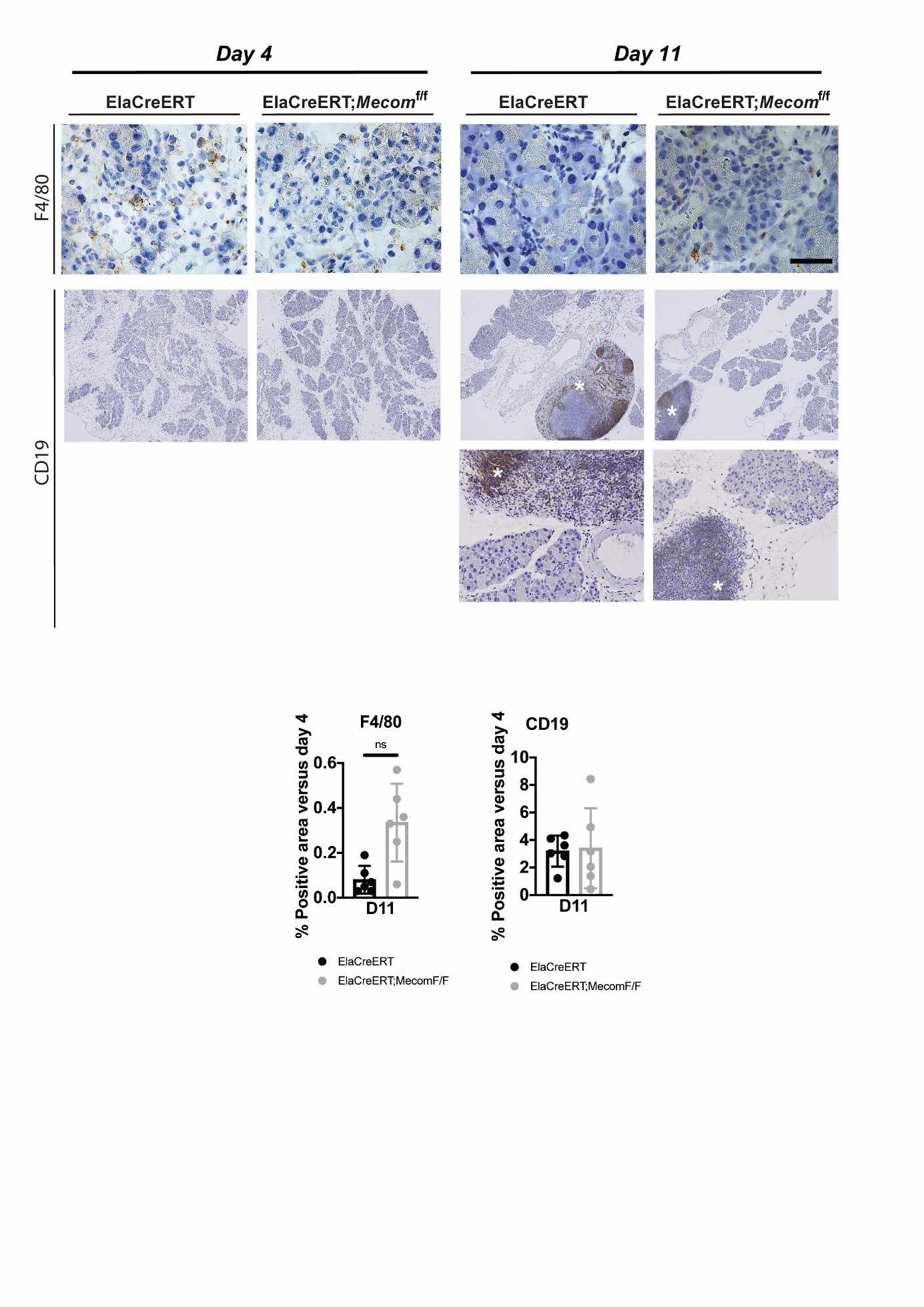

**SUPPLEMENTARY MATERIALS AND METHODS**

**List of primers**

| mEvi1 exon 3 fw | GCTATGATCAGCACAACCTTGTTG |
| --- | --- |
| mEvi1 exon 3 rv | TGTCTGCGACTACTCGGTAGAATATC |
| mHprt fw | GGCCAGACTTTGTTGGATTTG |
| mHprt rv | TGCGCTCATCTTAGGCTTTGT |
| mSox9 fw | TCGGTGAAGAACGGACAAGC |
| mSox9 rv | TGAGATTGCCCAGAGTGCTCG |
| hEVI1 exon 3 fw | CGAAGACTATCCCCATGAAACTATG |
| hEVI1 exon 3 rv | TCACAGTCTTCGCAGCGATATT |
| hHPRT1 fw | GGCTCCGTTATGGCGACCC |
| hHPRT1 rv | TGTGATGGCCTCCCATCTCCTT |
| hSOX9 fw | GAGGAAGTCGGTGAAGAACG |
| hSOX9 rv | ATCGAAGGTCTCGATGTTGG |
| hGAPDH fw | TGCACCACCAACTGCTTAGC |
| hGAPDH rv | GGCATGGACTGTGGTCATGAG |
